# The Role of Spatial Separation on Selective and Distributed Attention to Speech

**DOI:** 10.1101/2020.01.27.920785

**Authors:** Danna Pinto, Galit Agmon, Elana Zion Golumbic

**Author notes:** Corresponding Author: Dr. Elana Zion Golumbic, The Gonda Brain Research Center, Building 901, Bar Ilan University, Ramat Gan, Israel 5290002.

## Abstract

Processing speech in multi-speaker environments poses substantial challenges to the human perceptual and attention system. Moreover, different contexts may require employing different listening strategies. For instance, in some cases individuals pay attention *Selectively* to one speaker and attempt to ignore all other task-irrelevant sounds, whereas other contexts may require listeners to *Distribute* their attention among several speakers. Spatial and spectral acoustic cues both play an important role in assisting listeners to segregate concurrent speakers. However, how these cues interact with varying demands for allocating top-down attention is less clear. In the current study, we test and compare how spatial cues are utilized to benefit performance on these different types of attentional tasks. To this end, participants listened to a concoction of two or four speakers, presented either as emanating from different locations in space or with no spatial separation. In separate trials, participants were required to employ different listening strategies, and detect a target-word spoken either by one pre-defined speaker (Selective Attention) or spoken by any of the speakers (Distributed Attention). Results indicate that the presence of spatial cues improved performance, particularly in the two-speaker condition, which is in line with the important role of spatial cues in stream segregation. However, spatial cues provided similar benefits to performance under Selective and Distributed attention. This pattern suggests that despite the advantage of spatial cues for stream segregation, they were nonetheless insufficient for directing a more focused ‘attentional spotlight’ towards the location of a designated speaker in the Selective attention condition.

## 1. Introduction

Allocating attention appropriately in multi-speaker environments can be quite challenging, and yet is critical for behavior in many real-life situations (Bronkhorst, 2015, 2000; Ericson et al., 2004; Kaya and Elhilali, 2017; Shinn-Cunningham, 2008; Zion Golumbic et al., 2013). Performance is affected by the severity of the acoustic load imposed, but also by specific task demands and listener’s goals. For example, in some situations listeners attempt to pay attention to only one speaker and ignore all other sounds (“Selective Attention”), whereas other contexts require listeners to distribute their attention among several sound sources and monitor them all for potentially relevant information (“Distributed Attention”). Implementing either of these listening strategies in multi-speaker context arguably relies first on segregating the mixture into distinct speech streams (Bregman, 1990; Yabe et al., 2001) which is considered a pre-attentive process (Sussman, 2017; Symonds et al., 2019). Top-down attention can then be applied to the segregated streams to decide and prioritize which portion (or portions) of the acoustic scene will be further processed at a linguistic level (Best et al., 2010, 2006; Broadbent, 1958; Cherry, 1953), in accordance with behavioral goals.

Stream segregation depends critically on the distinctiveness of spectro-temporal features of different speakers (Bronkhorst, 2015; Elhilali et al., 2009; Moore and Gockel, 2002; Shamma et al., 2011; Thakur et al., 2015). For example, it is more difficult to segregate two recordings spoken by the same voice vs. those spoken by different speakers (Brungart et al. 2001; Ericson et al. 2004), and segregation of temporally asynchronous utterances is easier than that of synchronous speech (Humes et al., 2017). Another dimension that is particularly relevant for segregating speakers in real-world contexts is spatial separation, since competing speech sources typically do not emanate from the same location. Introducing spatial cues, in the form of interaural time or loudness differences (ITD/ILD respectively), can further improve stream segregation, a phenomenon referred to as ‘spatial release from masking’ (Ellinger et al., 2017; Eramudugolla et al., 2008; Maddox et al., 2012; Marrone et al., 2008).

As mentioned above, stream segregation is a pre-requisite for selecting and focusing attention on task-relevant speakers. However, some of the acoustic cues used for stream segregation may also be useful for directing attention in a top-down manner toward task-relevant portions of the scene and prioritize stimulus processing. In the case of both spectral and temporal cues, there is evidence that these cues are used to actively enhance sensitivity within specific neuronal populations in auditory cortex that encode the spectral features of attended sounds (Carlyon, 2004; Carlyon et al., 2001; Cusack et al., 2004; Fritz et al., 2007) as well as their expected timing (Lakatos et al., 2019; Nobre and Ede, 2017). Hence, these cues seem to be used both for stream segregation and for directing attention itself. However, as opposed to the spectro-temporal features, spatial encoding is not a primary dimension in the auditory system and is carried out by secondary computations (Keating and King, 2015; Kong et al., 2014; Shiell et al., 2018; Smith et al., 2010; Traer and McDermott, 2016). Therefore, it is more difficult to determine whether attention itself utilizes spatial information to prioritize processing of attended sounds, above and beyond its contribution to stream segregation. Although several studies have shown that performance on a variety of attention tasks is improved by increasing the spatial separation between attended and unattended speech (Best et al., 2006; Brungart, 2005; Gygi and Shafiro, 2014; Humes et al., 2006; Ihlefeld and Shinn-Cunningham, 2008a, 2008b, 2008c; Kidd et al., 2005), in these studies it is difficult to disentangle whether these effects are due to improved stream segregation for spatial speech, or to the effects of attention per se.

The current study was aimed at studying the potential dissociation between utilizing spatial cues for stream segregation and engagement of spatial attention in multi-speaker contexts. To do so, we take advantage of the different listening strategies required for Selective vs. Distributed attention. Although both require similar stream-segregation, only Selective attention involves prioritization of one speaker over all others. Therefore, if spatial separation between speakers also facilitates allocation of spatial attention, this is expected to benefit Selective attention more that Distributed attention. This is because in the case of Selective attention, an attention ‘spotlight’ can be directed towards a single to-be-attended location (Buschman and Miller, 2010; Posner, 1980), whereas under Distributed attention all locations remain task-relevant and no specific location can be prioritized for processing. Alternatively, if similar benefits for spatial separation are found in both Selective and Distributed attention tasks, this would suggest that the benefit is mostly derived from stream segregation itself and does not additionally reflect the engagement of spatial attention.

To this end, we compare performance on a target-word detection task in two attentional conditions. In the Selective attention condition, a single speaker (among a mixture) is designated to be monitored for target-words, whereas in the Distributed attention condition the target-word could be spoken by any of several concurrent speakers. We manipulated the number of concurrent speakers between two and four, since the fidelity of stream segregation and the utility of spatial cues has been shown to differ dramatically as the number of speakers increases (Brungart et al., 2001; Fairnie et al., 2016; Humes et al., 2017; Rosen et al., 2013; Simpson and Cooke, 2005). Critically, this design ensured that the acoustic load was equated between the two attention conditions, and did not impose additional working-memory demands, which is sometimes the case in studies comparing selective and divided attention (Best et al., 2010; Gygi and Shafiro, 2012; Humes et al., 2006; Koelewijn et al., 2014).

## 2. Materials and Methods

### 2.1 Participants

Participants were randomly assigned to two experimental groups, where they performed either a Spatial or Non-Spatial version of the same task. The study included a total of 57 participants, 27 in the Spatial condition (16 female; mean age 23.3 ± 2.3) and 30 in the Non-Spatial condition (21 female; mean age 23.5 ± 2.3). All participants were fluent Hebrew speakers, with self-reported normal hearing and no history of psychiatric or neurological disorders. Participants were paid or received course credit for participation. The study was approved by the Ethics Committee of Bar-Ilan University, and all participants read and signed an informed consent form prior to the commencement of the experiment.

### 2.2 Stimuli

The stimuli set consisted of 16 Hebrew mono-syllabic words recorded in four different voices (two male, two female; mean word duration=393 ms, SD=25; no significant difference in mean word-length between speakers, p>0.6). Audio editing of the individual words and their combination into sequences were performed using Audacity (www.audacityteam.org) and Matlab (The MathWorks, Inc). The perceived loudness of each speaker was equated offline and verified using inter-rater reliability testing. Individual words were concatenated to form 15 second-long sequences for each speaker (SOA between words randomly selected between 0.6 to 1.4 seconds; Figure 1A). The sequences of individual speakers were combined to create 2-speaker and 4-speaker conditions, and the timing of individual words was controlled to minimize temporal overlap between the words (2-speaker condition: maximal temporal overlap of 200 ms; 4-speaker condition: no more than 2 overlapping speakers, maximal temporal overlap of 250 ms).

**Figure 1.**
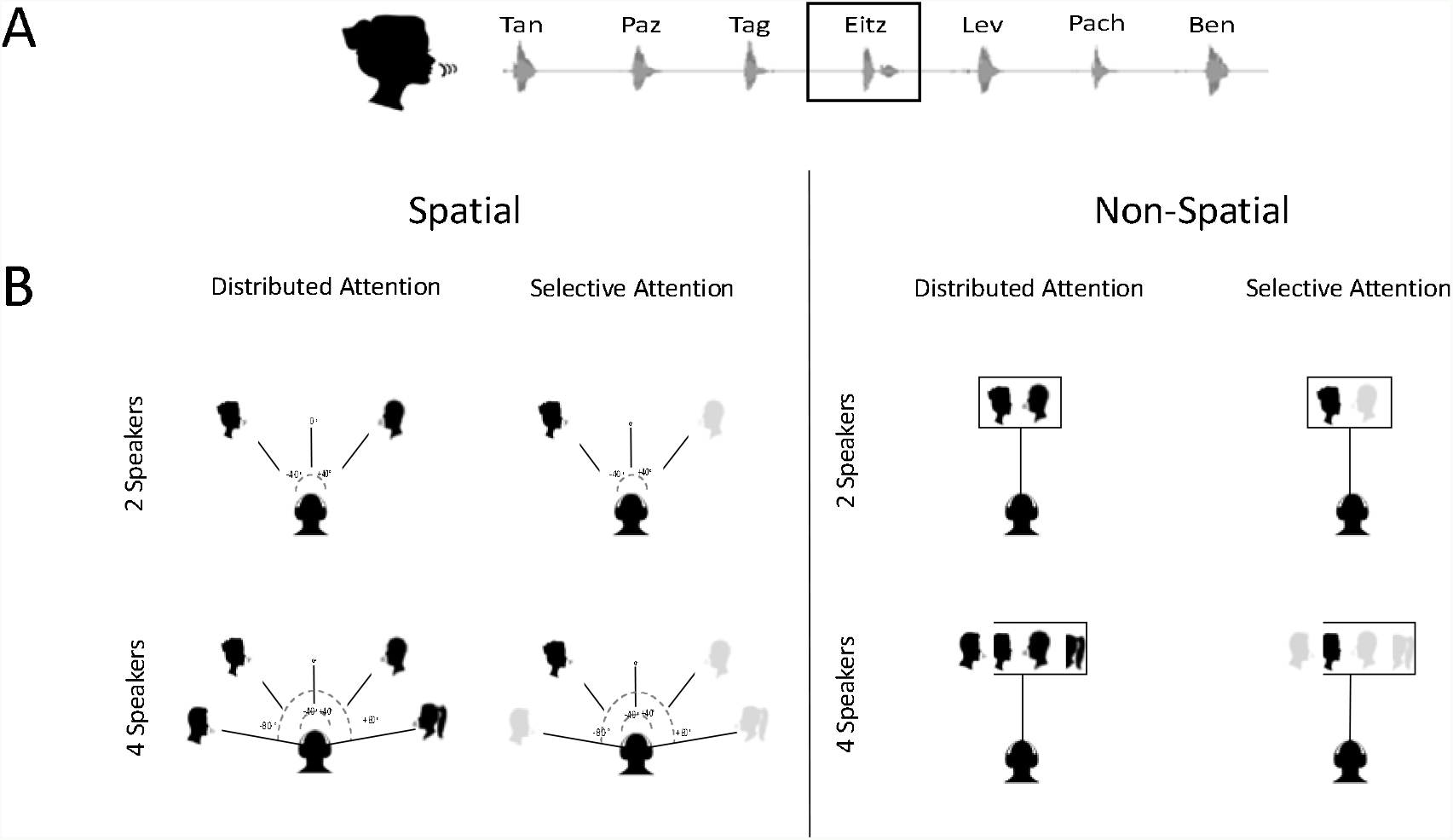
A. Example of the audio of a single speech stream, comprised of a sequence of mono-syllabic Hebrew words. The target word *Etz (tree)* is indicated with a gray square. B. Illustration of the experimental design: Spatiality was manipulated as a between-group factor, with the audio of concurrent speaker presented either diotically (Non-Spatial condition), or as originating from different locations (Spatial condition). The number of speakers (2 vs. 4) and attention type (Selective vs. Distributed) was manipulated within group. The color of the face-icons illustrates which speakers should be attended to (black) and which should be ignored (gray) in each condition.

In the Non-Spatial condition, the audio of all speakers was presented diotically with no spatial separation between them. In the Spatial condition, the audio of each speaker was convolved with head-related transfer functions (HRTF; KEMAR database, https://sound.media.mit.edu/resources/KEMAR.html) to create the perception of originating from different spatial locations. We used two location on the right (+40° and +80°) and two locations on the left (−40° and −80°; Figure 1B). All four locations were used in the 4-speaker condition, and in the 2-speaker condition one left-size and one right-side location were randomly selected in each trial. The attribution of speaker-to-location was randomized across trials.

### 2.3 Experimental Procedure

Participants performed a target-detection task under two condition, probing either Selective and Distributed Attention to speech (Figure 1B). The Selective Attention condition, required participants to attend to one designated (“attended”) speaker and count the occurrences of a target-word uttered only by this speaker. The identity of the “attended” speaker was randomly changed in each block, and participants were familiarized with their voice prior to each block. In the Distributed Attention condition, participants were instructed to count how many times the target word was spoken by <u>any</u> of the concurrently presented streams. The same target word (tree; “*Etz*”) was used in all conditions and could occur either 0,1 or 2 times per trial. Participants were asked to give their response at the end of each trial. Both the Selective and Distributed tasks were performed at two difficulty levels, with either 2-or 4-concurrent speakers. Since the similar stimuli were used in the two tasks, perfectly matched for both low-level acoustics and working-memory load, any differences in performance are attributed to the different type of attention required.

Selective and Distributed trials were presented in 16 separate blocks (8 per condition), with order counter-balanced order across participants. Each block consisted of both 2-speaker and 4-speaker trials (4 of each per block). This yielded a total of 32 trials per Attention-Type (Selective/Distributed) × Number of Speakers (2/4) combination. A third between-subject variable was Spatial Separation.

The experiment was conducted in a sound-attenuated booth, and speech stimuli were presented through headphones (Sennheiser, HD 280 Pro). The experiment was programmed using Psychopy software (Peirce et al., 2019), and responses were recorded using a response box (Cedrus RB 840). After the experiment participants completed two additional tasks for assessing non-verbal general-intelligence functioning (g-factor Test of Nonverbal Intelligence 3rd Edition; TONI-3) and working memory capacity (WMC, Wechsler Adult Intelligence Scale-Fourth Edition).

### 2.4 Statistical Analysis

Single trials were categorized as “correct” if participants indicated the correct number of occurrences of the target-word. To analyze the differences in accuracy between conditions, single trials were fit using a generalized linear mixed-effects regression model with a logit link function, as implemented in R’s glmer function (lme4 package) (Bates et al. 2015). A mixed-effect model accounts for variability across subjects and correlations within the data (Baayen, Davidson & Bates, 2008). A logit-based model allows to accurately analyze binomial data in repeated-measures designs (Jaeger, 2008).

The model included two within-subject factors (Attention Type and Number of Speakers) and one between-subject factor (Spatial Separation), as well as all of their interactions. We also included individual scores on the working memory test as well as non-verbal IQ scores as fixed effects, and included random by-subject intercepts and random slopes for Attention Type. All factors were sum-coded, i.e. the coefficients represent deviation from the grand mean, thus allowing for orthogonal contrasts. The reported p-values are based on asymptotic Wald tests included in the summary of R's glmer function.

## 3. Results

We found significant main effects for all three experimental factors: Attention Type, Number of Speakers and Spatial Separation (Figure 2A and Table 1). Participants performed better in the Selective attention than in the Distributed attention condition (β=0.69, z=2.62, p<0.009), and performance was higher in the 2 speakers vs. the 4 speaker conditions (β= −1.46, z=8.09, p<10^−16^). In addition, accuracy was higher when audio was presented with Spatial Separation vs. the Non-Spatial condition (β= −1.00, z=3.72, p<0.0002). There was also a significant interaction between Spatial Separation × Number of Speakers (β=0.80, z=3.23, p<0.001), indicating that Spatial Separation improved performance on the 2-speaker condition more than in the 4-speaker condition. This was confirmed in follow-up analyses, where a significant effect for Spatial Separation was found in the 2-speaker condition (z=−2.034, p< 0.05), but not in the 4-speaker condition (z=−0.350, p>0.7). However, none of the interactions with Attention Type were significant (p>0.7), suggesting that Spatial Separation has a similar effect on both types of attention. None of the effects correlated significantly with individual cognitive traits as assess by WM capacity (p>0.6) or non-verbal IQ (p>0.1). In this sample, WM capacity and non-verbal IQ were also not correlated between themselves (Pearson’s r = 0.08).

**Table 1:** Summary of all results in the glmr model testing for effects of Attention Type, Number of Speakers and Spatial Separation, and their interactions.

| *Factor* | *Difference ± SE(%)* | *β* | *z* | *p* |
| --- | --- | --- | --- | --- |
| ***Spatial vs. Non Spatial*** | 3 ± 2% | 1.00 | 3.72 | 0.001 |
| ***Four vs. Two Speakers*** | -8 ± 1% | 1.46 | 8.09 | 6.22 x 10<sup>-16</sup> |
| ***Selective vs. Distributed*** | 6 ± 1% | 0.69 | 2.63 | 0.01 |
| ***Number of Speakers \* Attention Type*** | 6 ± 1% | 0.36 | 1.18 | 0.24 |
| ***Number of Speakers \* Spatiality*** | -2 ± 2% | 0.80 | 3.23 | 0.00 |
| ***Attention Type \* Spatiality*** | 0 ± 1% | -0.25 | -0.73 | 0.47 |
| ***Attention Type \* Spatiality \* Number of Speakers*** | 5 ± 3% | 0.07 | 0.17 | 0.86 |

**Table 2:** Summary of all results in the glmr model testing for effects of Attention Type, Number of Speakers and Spatial Separation, and their interactions.

| *Factor* | *Difference ± SE(%)* | *β* | *z* | *p* |
| --- | --- | --- | --- | --- |
| ***Spatial vs. Non Spatial*** | 3 ± 2% | 1.16 | 3.43 | 0.001 |
| ***Selective vs. Distributed*** | 4 ± 1% | 1.12 | 2.57 | 0.01 |
| ***Same Vs. Different Speaker Gender*** | -2 ± 1% | 1.05 | 3.30 | 0.00 |
| ***Speaker Gender\* Attention Type*** | 2 ± 2% | 0.10 | 0.19 | 0.85 |
| ***Speaker Gender \* Spatiality*** | -6 ± 2% | 1.15 | 2.78 | 0.01 |
| ***Attention Type \* Spatiality*** | -2 ± 2% | 0.08 | 0.16 | 0.88 |
| ***Attention Type \* Spatiality \* Speaker Gender*** | 0 ± 3% | 0.32 | 0.44 | 0.66 |

**Figure 2.**
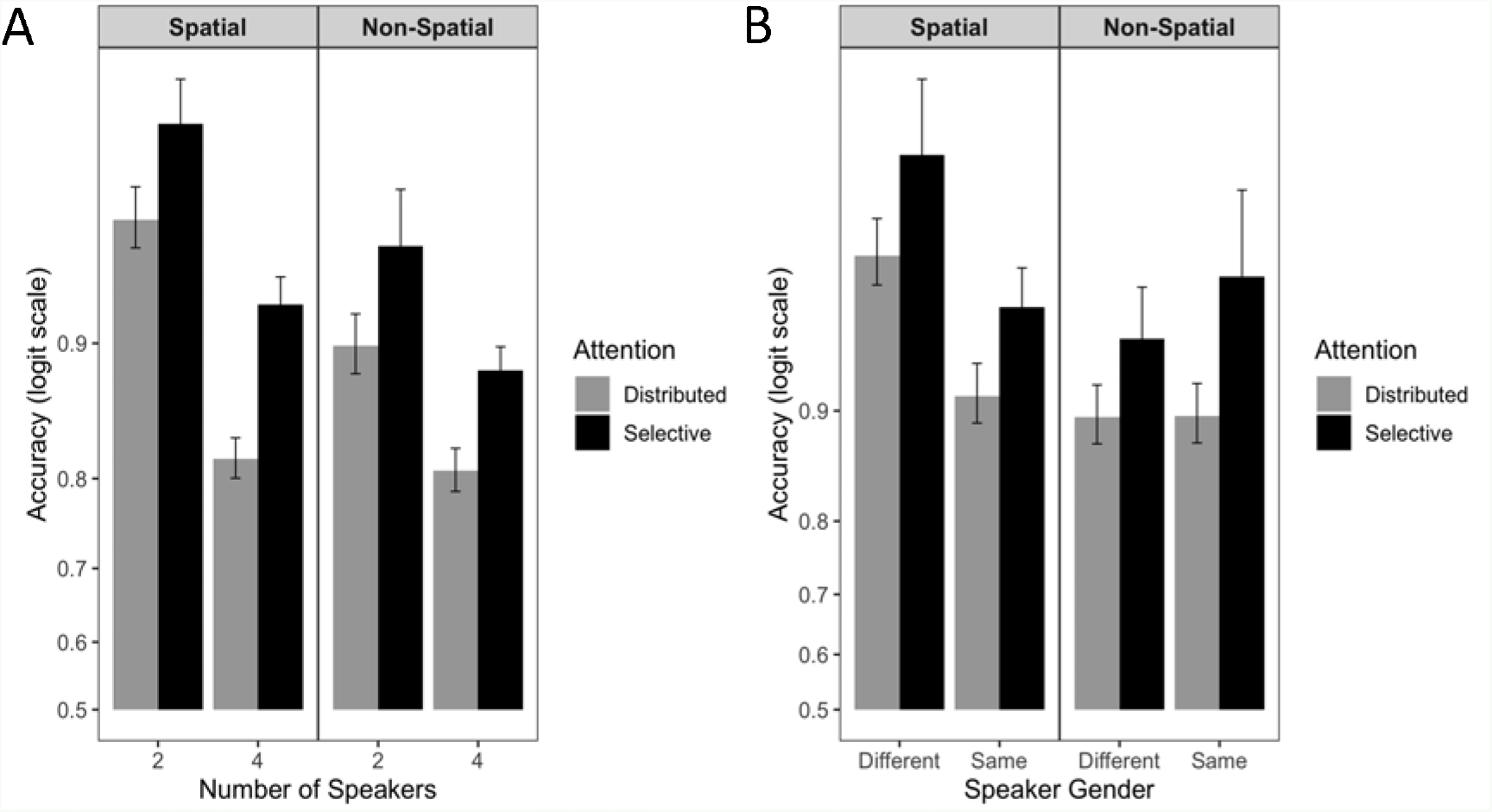
A. Performance accuracy across the three conditions (Spatial Separation, Attention Type, and Number of Speakers). B. Performance accuracy in the 2-speaker condition, separated according to whether they were of the same or different biological gender. Results in both panels are displayed on a logit scale.

A follow-up analysis focusing on the 2-speaker conditions was conducted, to test whether the acoustic similarity between speakers interacted with the type of attention and spatial separation. To this end, we separated trials in which the competing speakers were of the same biological gender (male/male and female/female), and those with speakers of different biological genders (female/male). A similar glmer model was constructed with 3 independent variables (Spatial Separation, Attention Type, and Speaker Gender Similarity). As expected from the main analysis, there were significant main effects for spatial separation (β= −1.15, z=3.428, p<0.0007) and attention type (β= −1.12, z=2.57, p<0.02). The main effect of speaker-gender similarity was also significant (β= −1.05, z=3.304, p<0.001), as accuracy in different-gender trials was significantly higher than same-gender trials. This effect also interacted significantly with spatial separation (β= 1.15, z=2.78, p<0.006), with the effect of speaker-gender similarity larger in the spatial vs. non-spatial condition. Follow up analyses confirmed that speaker-gender similarity did no significantly affect performance in the non-spatial condition (β= −0.198, z=0.83, p=0.41), and the effect was limited to the spatial condition (β= 1.107, z=4.041, p<0.0001). This pattern indicates that performance is optimal when the two competing voices are distinct from each other in both pitch and in space, however having distinct voices did not facilitate performance in the non-spatial condition. Importantly, here too there was no significant interaction with attention type, suggesting that here too acoustic-similarity affected performance on both tasks similarly.

## 4. Discussion

Here we studied the contribution of Spatial Separation between competing speakers to performance on two different types of attentional tasks: Selective vs. Distributed attention to speech. We found that Spatial Separation improved performance on both tasks, but only in the two-speaker condition. When four speakers were presented concurrently, spatial separation did not improve performance significantly. Moreover, pitch separation between two competing speakers (operationalized as same vs. different biological gender) also improved performance, but only when spatially separated. Critically, although Distributed attention is overall more difficult than Selective Attention, resulting in reduced performance in this study as well as previous work (Baldock et al., 2018; Best et al., 2006; Brungart, 2001; Koelewijn et al., 2014; Lambez et al., 2019), spatial separation affected performance on both tasks in a similar fashion, with no significant interaction between them. The lack of a difference between the effects of spatial cues on these two extremely different types of attentional tasks leads us to suggest that spatial cues play a limited role in directing top-down attention in multi-speaker environment, beyond their contribution to stream segregation. We elaborate on this interpretation in the following sections.

### 4.1 Contribution of Spatiality to Stream Segregation

Stream segregation is considered a pre-attentive process, that is applied to incoming sounds even if they are outside the focus of attention (Sussman, 2017; Symonds et al., 2019).

It is carried out based on the conjunction of multiple stimulus-features including pitch, timbre and temporal structure, with spatial cues such as ITD and ILD contributing as well albeit to a lesser degree (David et al., 2017; Snyder and Elhilali, 2017; Stainsby et al., 2011). Previous studies have shown that segregation of two competing speakers is relatively good, but that performance drops sharply as the number of concurrent speakers increase beyond two (Brungart et al., 2001; Humes et al., 2017; Rosen et al., 2013; Simpson and Cooke, 2005). This is true for spatial and non-spatial speech alike (Kawashima and Sato, 2015). However in the case of two speakers, spatial cues provide an additional advantage of spatial release from masking, that diminishes as the number of sources increases (Freyman et al., 2004; Yost, 2017; Yost et al., 2019). In other words, not only is segregation of four speakers using acoustic features more difficult, but adding spatial cues does not contribute substantially to their unmasking. The current results are consistent with previous findings, showing a decrease in performance between two to four speakers, and an interaction with spatiality indicating a larger advantage for spatial cues in the two-speaker condition.

Interestingly, we find that spatial separation also interacted with the effect of the competing speakers’ biological gender on performance. Previous studies have established that the characteristic differences in voice acoustics of female and male voices (Huber et al., 1999; Read, 1992) result in reduced performance on selective attention tasks among same-gender versus different-gender speakers, although results in distributed attention tasks are inconsistent (Brungart, 2001; Brungart et al., 2001; Darwin et al., 2003; Humes et al., 2006; Xia et al., 2015). The current results indicate that spatial and acoustic separation have additive beneficial effects for improving performance. However, no benefit for different-gender speakers was observed in the non-spatial condition, suggesting that optimal stream segregation among competing speakers requires a conjunction of spectral and spatial cues.

### 4.2 No Evidence for Engagement of Spatial Attention

Beyond the advantage of spatial and acoustic cues for stream segregation, in the current study we were further interested in understanding whether attention itself utilizes spatial information to direct top-down attention toward particular locations in space. If this is the case, we expected to find a larger advantage for spatial separation for Selective Attention, where the attention ‘spotlight’ could be directed specifically toward a particular location and other locations can be suppressed, as is commonly found in the visual domain (Falkner et al., 2010). At the same time, Distributed Attention should become more difficult in the spatial condition since it would require monitoring input from several locations rather than from one central location, which is expected to be more effortful (Best et al., 2006; Koelewijn et al., 2014). Counter to this original hypothesis, we found no difference in the way spatial cues affected the two types of attention. For two speakers, both Selective and Distributed attention benefitted from spatial cues, and for four speakers neither type of attention was affected by spatiality. Moreover, the advantage found for different-gender speakers in the two-speaker Spatial condition was not different for Selective and Distributed attention. Hence, the current results do not support the engagement of spatial attention in either task, beyond the (pre-attentive) contribution of spatial cues to stream segregation.

Several neuroimaging studies have pointed to a distinction between brain regions in the auditory system that utilize spatial information for the purpose of stream segregation and regions encoding the spatial location of an auditory source (Shiell et al., 2018; Smith et al., 2010). Localization of sounds in space is computationally challenging, particularly given the effects of reverberation in natural settings (Keating and King, 2015; Traer and McDermott, 2016), and likely relies on different underlying mechanisms than stream segregation (Snyder and Elhilali, 2017). The dissociation between utilization of spatial cues for the purpose of stream segregation and determining their specific spatial location is also supported by behavioral findings demonstrating increased sensitivity and discriminability for spatially segregated sounds, albeit with poor ability to report the spatial location of the target (Klatt et al., 2018; Middlebrooks, 2013; Weller et al., 2016). In other words, determining that two inputs originate from different locations is possible even when their exact location cannot be reliably discerned. However, engagement of **spatial attention** does depend on having a reliable representation for the spatial location of an attended stimulus (Kong et al., 2014). The lack of a difference between Selective and Distributed attention in the spatial condition, may suggest that, at least in this particular case, the spatial cues provided were insufficient for engaging spatial attention per se.

This is not to say that it is not possible to engage spatial auditory attention to enhance attention to speech. For example, when more detailed spatial information can be provided regarding the location of each speaker, as in the case of audio-visual speech (Blackburn et al., 2019) or spatial cueing, spatial attention might be engaged. Indeed, (Kopco et al., 2010) demonstrated that giving participants information regarding the location of potential maskers assisted selective-attention performance, although the benefit depended on the specific spatial configuration (see also Jones and Litovsky 2008; McCloy and Lee 2015). Similarly, (Ihlefeld and Shinn-Cunningham, 2008a) report an advantage of spatial separation on selective attention performance, but only when participants were explicitly told to attend to a particular location, whereas when attention was directed towards an acoustic feature, no advantage was observed for spatial separation. (Singh et al., 2008) found that selective attention improved when participants could anticipate where the attended speaker would occur, further supporting an added cognitive value for spatial separation. Together with the lack of a benefit for spatial cues in the Selective attention condition in the current study, where spatial information was stimulus-driven and implicit, these findings suggests that spatial attention is not automatically engaged just because spatial cues are available. These results invite future follow up research in order to fully characterize the conditions under which top-down spatial attention is utilized to improve performance in multi-speaker contexts.

## Acknowledgements

This work was supported by the Israel Science Foundation Center of Excellence grant 51/11 to ezg and the Bi-national Science Foundation Grant #2015385 to ezg.

## Data Statement

The full data set is openly available on Mendeley Data doi: 10.17632/29yssmwmgp.2

